# Cross-reactivity of r*Pvs*48/45, a recombinant *Plasmodium vivax* protein, with sera from *Plasmodium falciparum* endemic areas of Africa

**DOI:** 10.1101/2024.04.10.588966

**Authors:** Saidou Balam, Kazutoyo Miura, Imen Ayadi, Drissa Konaté, Nathan C. Incandela, Valentina Agnolon, Merepen A Guindo, Seidina A.S. Diakité, Sope Olugbile, Issa Nebie, Sonia M Herrera, Carole Long, Andrey V. Kajava, Mahamadou Diakité, Giampietro Corradin, Socrates Herrera, Myriam Arevalo Herrera

**Affiliations:** International Center for Excellence in Research (ICER-Mali), University of Sciences, Techniques and Technologies of Bamako (USTTB), Bamako, Mali; Laboratory of Malaria and Vector Research, National Institute of Allergy and Infectious Diseases, National Institutes of Health, Rockville, MD 20852, USA; Immunobiology Department, University of Lausanne, Lausanne, Switzerland; Division of Immunology and Allergy, Centre Hospitalier Universitaire Vaudois (CHUV), Lausanne, Switzerland; Groupe de Recherche Action Santé (GRAS), Burkina Faso, West Africa; Caucaseco Scientific Research Center, Cali, Colombia; Montpellier Cell Biology Research Center (CRBM), University of Montpellier, CNRS, France; Malaria Vaccine and Drug Development Center, Cali, Colombia

**Keywords:** r*Pvs*48/45 protein, *P. falciparum* African sera, Cross-reactive antibodies, Transmission-blocking vaccine

## Abstract

**Background:** *Ps*48/45, a *Plasmodium* gametocyte surface protein, is a promising candidate for malaria transmission-blocking (TB) vaccine. Due to its relevance for a multispecies vaccine, we explored the cross-reactivity and TB activity of a recombinant *P. vivax Ps*48/45 protein (r*Pvs*48/45) with sera from *P. falciparum*-exposed African donors.

**Methods:** r*Pvs*48/45 was produced in Chinese hamster ovary cell lines and tested by ELISA for its cross-reactivity with sera from Burkina Faso, Tanzania, Mali, and Nigeria – In addition, BALB/c mice were immunized with the r*Pvs*48/45 protein formulated in Montanide ISA-51 and inoculated with a crude extract of *P. falciparum* NF-54 gametocytes to evaluate the parasite-boosting effect on r*Pvs*48/45 antibody titers. Specific anti-*rPvs*48/45 IgG purified from African sera was used to evaluate the *ex vivo* TB activity on *P. falciparum,* using standard mosquito membrane feeding assays (SMFA).

**Results:** r*Pvs*48/45 protein showed cross-reactivity with sera of individuals from all four African countries, in proportions ranging from 94% (Tanzania) to 40% (Nigeria). Also, the level of cross-reactive antibodies varied significantly between countries (p<0.0001), with a higher antibody level in Mali and the lowest in Nigeria. In addition, antibody levels were higher in adults (≥ 17 years) than young children (≤ 5 years) in both Mali and Tanzania, with a higher proportion of responders in adults (90%) than in children (61%) (p<0.0001) in Mali, where male (75%) and female (80%) displayed similar antibody responses. Furthermore, immunization of mice with *P. falciparum* gametocytes boosted anti-*Pvs*48/45 antibody responses, recognizing *P. falciparum* gametocytes in indirect immunofluorescence antibody test. Notably, r*Pvs*48/45 affinity-purified African IgG exhibited a TB activity of 61% against *P. falciparum* in SMFA.

**Conclusion:** African sera (exposed only to *P. falciparum)* cross-recognized the r*Pvs*48/45 protein. This, together with the functional activity of IgG, warrants further studies for the potential development of a *P. vivax* and *P. falciparum* cross-protective TB vaccine.

## 1. INTRODUCTION

Malaria is a human parasitic disease caused by *Plasmodium* species, including *Plasmodium falciparum, P. vivax, P. malariae, P. ovale* and *P. knowlesi* [1]. Currently, particular attention is given to controlling *P. falciparum* and *P. viva*x, responsible for over 90% of the 247 million malaria cases reported globally in 2021. Approximately 93% of these cases occurred in Africa, where *P. falciparum* causes 99% of clinical cases[2, 3]. In contrast, both species are endemic in Asia and America, where *P. vivax* accounts for ∼74% of malaria cases[2, 3].

As a result of improved access to first-line treatment, rapid diagnostics tests, and vector control measures, a significant decrease (∼54%) in malaria cases was observed worldwide between 2000 and 2015, stimulating malaria eradication efforts [2–4]. However, certain regions have shown a significant increase in malaria cases during the last few years [5], constituting an important obstacle to this goal. This appears to be due to the emergence of resistance to first-line anti-malarial drugs like artemisinin derivatives^2^ [6–9] and mosquito resistance to insecticides, both of which appear to be rapidly disseminating across the globe [10–13]. Furthermore, the vast presence of asymptomatic and sub-microscopic infections also contributes to malaria transmission levels [14–16]. Therefore, efforts must be intensified to develop novel tools and strategies, such as vaccines, to strengthen malaria control and elimination efforts.

Vaccines are considered a highly cost-effective method for combating infectious diseases and are garnering increasing interest within the malaria public health agenda [17–21]. Following extensive evaluations in a phase III clinical trial with children across several African countries, [21–29], *P. falciparum* RTS, S and R21, based on the circumsporozoite (CS) protein fused to the Hepatitis B surface antigen (HBsAg) and expressed in yeast, have emerged. RTS,S is the first malaria vaccine to be recommended by the World Health Organization (WHO) for widespread use among children in sub-Saharan Africa, and other regions which display moderate to high *P. falciparum* transmission rates[30, 31]. However, RTS,S/AS01 exhibits modest efficacy, 39% against clinical malaria and 29% against severe malaria during a median of 48 months follow-up period [21]. Nevertheless, other *P. falciparum* vaccine candidates like R21/Matrix-M has exhibited a remarkable protective efficacy of 75% in children living in areas with high malaria seasonal transmission rates, proving to be safe, highly immunogenic, and promising in terms of efficacy [32–34]. Additionally, a *Pf*SPZ candidate based on whole cryopreserved parasites is also being evaluated in clinical trials in African countries [32, 35–38].

Although *P. vivax* vaccine research lags behind *P. falciparum*, several vaccine-candidate antigens from different parasite development stages are also being investigated in preclinical and clinical phases. A *P. vivax* CS synthetic vaccine formulation has reached phase 1 and 2 clinical evaluations [39–41]. At the same time, antigens like *Pvs*48/45, *Pvs*25, and *Pvs*230, which are expressed in sexual parasite forms [42–49], as well as several antigens from asexual parasite stages containing coiled-coil motifs [50, 51], are among the most relevant *P. vivax* vaccine candidates [50, 52–54].

*Pvs*48/45 is an orthologous protein to *P. falciparum* and is currently being investigated as a TB vaccine candidate. Previous studies have shown that individuals from malaria-endemic areas harbor antibodies specific to this protein, with *ex-vivo* TB activity [55, 56]. Generally, *Pv*s48/45 genes are highly conserved and display an overall sequence homology of ∼56% between *P. falciparum* and *P. vivax* [57–59]. Furthermore, the low genetic polymorphism does not appear to influence the tertiary structure or the antigenic cross-reactivity [60]. Importantly, the functional analysis of the Ps*48/45* gene in *P. berghei* established its crucial role in the fertility of male gametes [47, 48, 58].

In the case of *P. vivax,* recent studies have shown that communities endemic with *P. vivax* or *P. falciparum* transmission exhibit high recognition of the *Pvs*48/45 protein and demonstrate TB activity in *ex vivo* DMFA [61, 62]. In these studies, human samples were collected from both *P. vivax* and *P. falciparum* endemic areas, so cross-reactivity in humans could not be studied. Furthermore, in mouse immunization studies, it had been observed that mice immunized with *Pvs*48/45 displayed strong cross-reactive antibodies to *Pfs*48/45 and showed that *Pfs*48/45 and *Pvs*48/45 antigens were able to cross-boost each other in cross-boosting experiments [55, 63]. In this study, we used sera from Africa where only *P. falciparum* was present to check the natural antibody cross-reactivity and TB activity in humans.

## 2. MATERIALS AND METHODS

### 2.1 Ethics, consent, and permissions

This study was conducted as part of a research protocol on malaria immunity in Mali (Protocol # 08-I-N120). All serum samples were anonymized archived and stored at -80°C, and the same samples used in our previous or collaborative studies [64–69]. Samples from Burkina Faso (BF) were collected in 1998 in the village of Goundry, 30km from Ouagadougou (the capital city of Burkina Faso. In 1998, no authorization was required for research study in BF. For Mali (ML), samples were collected from 2009 to 2011 in Kenieroba village located in Bancoumana district, 73 km from Bamako (the capital city), and in Dangassa village in Kourouba town, 80 km from Bamako. Approval was obtained from the Ethical committee (EC) of Faculty of Medicine, Pharmacology and Odonto-Stomatology (FMPOS), University of Bamako, Mali (0840/FMPOS) . For Tanzania (TZ), samples were selected from those collected from 1982 to 1984 during a large-scale community-based study undertaken in Ifakara village in the Kilombero District in Morogoro. The authorization was obtained from the Commission for Science and Technology (UTAFITI NSR/RCA 90). Samples from Nigeria (NIG) were collected on March 2^nd^ of 2007, from donors living in Lagos, southwest, Ethical approval was obtained from the Lagos State University Teaching Hospital (LASUTH) ethical review committee [70]. Sera from healthy Swiss adults were from those who gave their informed consent (IC) to participate in malaria vaccine research in 2012 (code: NCT01605786de a). Written IC for collection of iRBCs for IFAT and sera for ELISA were obtained from all adults. Informed assent (IA) was obtained from children in addition to IC from their parents or legal guardians. The animal studies were approved by the Research Ethics Committee of the School of Health, Universidad del Valle (Cali-Colombia) (Code: 031-015).

### 2.2 Blood samples

Blood samples were collected from adults (≥17-year-old) and children (≤5-year-old) donors living in four different malaria-endemic countries of Africa: Burkina Faso (BF; adults N=35), Mali (adults N=62 and children N=97), Tanzania (TZ; adults N=83 and children N=63) and from Nigeria (NIG; adults N=10). Whole blood was collected in EDTA tubes, and then the sera were extracted by centrifugation and stored at -80°C prior to the different tests. Samples from Burkina Faso (BF) and Nigeria (NIG) were collected from donors living in urban settings. In contrast, samples from TZ and Mali were obtained from rural settings where exposure to malaria is potentially higher than in urban settings. In BF, samples were collected in 1998 from Ouagadougou (the capital city), and those of Mali were collected from 2009 to 2011 from Kenieroba, Bozokin, and Fourda, villages located in the Bancoumana district, about 55 km from Bamako (the capital city), and Dangassa, a village in the Kourouba District, 80 km from Bamako. Samples from Tanzania (TZ) were collected from 1982 to 1984 during a large-scale community-based study undertaken in Ifakara (village in the Kilombero District in Morogoro). Samples from NIG were obtained from adult donors living in Lagos city, southwest NIG. Anonymized sera from 10 healthy Swiss adults non-exposed to malaria (who had no malaria history) who gave their ICs to participate in malaria vaccine research (2012, study NCT01605786) were used as negative controls.

### 2.3 BALB/c mice immunization and bleeding

In a previous study [55], the immunogenicity of CHO-r*Pvs*48/45 protein emulsified in Montanide ISA-51 was evaluated in twelve male and female, 6-8 weeks old BALB/c mice (six experimental and six control). After three doses of 20 μg (subcutaneously, s.c.) of the protein (days 0, 20, and 40), all animals of the experimental group seroconverted and reached ELISA titers up to 1:10^6^ [55]. Control mice were immunized with saline solution formulated in Montanide ISA 51. In this study, when specific anti-CHO r*Pvs*48/45 antibodies had waned to baseline (day 260), a lysate of 5x10^5^ extract of mix gametocytes (∼80% mature forms) from *P. falciparum* NF54 parasite cultures were formulated in Montanide ISA 51 and inoculated intramuscularly (i.m.) to assess the potential boosting effect of *P. falciparum* gametocytes on anti-CHO r*Pvs*48/45 antibody titers. Four weeks after, mice were bled from the submandibular veins (∼100 μL), and specific anti-*rPvs*48/45 antibodies were analyzed by ELISA.

### 2.4 *Pvs*48/45 protein sequence and its homology analysis with *Pfs*48/45

The primary sequence of *Pvs*48/45 presented 6-Cys domains with 15 Cys residues, the N-terminal signal peptide, and the C-terminal GPI anchor predicted by the Signal P 3.0 and the GPI-SOM servers. Sequences of the *Pvs*48/45 (PlasmoDB accession number ALS19583.1) and *Pfs*48/45 (PlasmoDB accession number CAA57308.1) orthologous proteins were determined using the Salvador I genome database (PlasmoDB). The two sequences were then compared for homology using Blastp (protein-protein BLAST) (Suppl. figure 1) [57, 59].

### 2.5 Recombinant CHO-r*Pvs*48/45 protein production, purification, and analysis

The full-length CHO-*rPvs*48/45 protein was produced by Transient Gene Expression (Excellgene SA, Monthey, Switzerland) with sufficient viable cell culture biomass of suspension-adapted CHO-Express™ cells as described before [62, 71]. Briefly, the full-length *Pvs48/45* codon harmonized gene was produced in CHO cell lines. All production cultures (post-transfection) were performed in serum-free, animal-protein-free medium (low protein content). A total of 150 mg of this antigen was produced after a single purification step on IMAC-FPLC. Protein identity was confirmed using SDS-PAGE analysis of CHO-r*Pvs*48/45 protein under reducing (0.05 mol/L dithiothreitol) and non-reducing conditions together with immunoblot and mass spectrometry (LC-MS/MS) (Suppl. figure 2) [55].

### 2.6 ELISA assays

***Indirect ELISA*** was performed using Maxisorp 96-well microtiter plates (Thermo Scientific, Ref. 442404) coated with 50 µL of a 2 µg/mL solution of the r*Pvs*48/45 protein solution overnight at 4°C. The plates were incubated (blocked) for 1 hour at RT with phosphate buffered saline (PBS) containing 3% non-fat milk powder (PBSx1-milk 3%), then incubated for 2 hours at room temperature (RT) with human sera at a dilution of 1:200 in PBS containing 3% milk and 0.05% Tween 20 (PBS-T). The plates were then washed four times with PBS-T. Goat anti-human IgG conjugated with horseradish peroxidase (HRP) was used as the secondary antibody at a dilution of 1:2000 (Life technologies, Ref H10307) in PBS-T-milk for 1 hour at RT. After four time washing with PBS-T, the signals were revealed using TMB substrate reagent (BD OptEIA, cat 555214) for 25 min in the dark at RT followed by a second stage blocking using 1M sulphuric acid (Merck, 1.00731.1000). Optical density (OD) was measured at 450 nm/630 nm using a TECAN Nano Quant Infinit M200 PRO spectrophotometer. ELISA was considered positive when a sample’s ODs was higher than the mean OD + 3 SD of negative controls (naïve human sera, NHS from Swiss naïve donors) diluted to 1:200.

***Competition ELISA*** was performed by incubating sera from BF (BF40 and BF70) at a dilution of 1:200 with r*Pvs*48/45 at 10-fold serial dilutions from starting at 300 µg/mL for 1 hour at RT prior to transferring the mixture to wells coated with the same r*Pvs*48/45. Plates were then incubated for 30 min at RT, and the reactivity was determined as previously described [72]. Each test was performed in duplicate. The percentage of inhibition was thus calculated as: (mean of Ab OD with competitor protein /mean antibody OD without competitor protein) x100.

### 2.7 Affinity purification of anti-*Pvs*48/45 antibodies

A serum pool collected from TZ adult donors with high anti-CHO-r*Pvs*48/45 antibody titers was used for IgG purification, as described previously [72, 73]. To prepare the antigen-Sepharose conjugate, CNBr-sepharose 4B (Amersham Bioscience AB, Uppsala, Sweden) was activated with 1 mM HCl. Then, 5 mg of the CHO-r*Pvs*48/45 protein was dissolved in 1 mL of coupling buffer (0.1 M NaHCO3 containing 0.5 M NaCl, pH 8.0). The sera were diluted 5-fold with PBS (1x) containing 0.5 M sodium chloride and were mixed with the antigen-sepharose conjugate and stirred O/N gently at 4°C. The antigen-sepharose beads were then washed with 5 mL of TRIS (20 mM containing 0.5 M NaCl, pH 8.0) and then with 5 mL of TRIS (20 mM, pH 8.0). The bound antibody was eluted with a solution containing glycine (0.1 M, pH 2.5). The fractions (F1, F2, F3) were collected in TRIS solution (1 M, pH 8.0) to instantly neutralize the solutions before dialyzing them against phosphate buffer (0.1M, pH 7.0). The antibody (IgG) concentration of each fraction was determined by the absorbance of the solution at 280 nm [72, 73]. In addition, ELISA was used to determine the recognition of r*Pvs*48/45 by each of the purified antibody fractions.

### 2.8 Indirect Immunofluorescence Antibody Test (IFAT)

Cross-recognition of *P. vivax* and *P. falciparum* was determined by IFAT using sera from mice immunized with CHO-r*Pvs*48/45. To this end, *P. vivax*-infected red blood cells (iRBC) were obtained from infected patients (code CECIV 1506-2017). White blood cells were separated using a 45% percoll gradient centrifugation at 5,000 rpm for 10 min, and an enriched *P. vivax* gametocytes fraction was obtained [74] and used to prepare 12-well glass microscope slides. For *P. falciparum,* mature gametocytes were obtained from *in vitro* culturing of the *Pf*-NF-54 parasite isolate, which were used to prepare IFAT glass microscope slides as described before [75]. IFAT slides were kept at -70°C prior to use. For IFAT reaction, slides were incubated with a pool of serum samples (at a 1:200 dilution) obtained from mice immunized with r*Pvs*48/45 and with a pool of control sera from naïve mice (at a 1:20 dilution) in PBS-Evans blue for 30 min. After PBS washing, slides were incubated with fluorescein isothiocyanate (FITC) conjugated anti-mouse IgG antibody at 1:100 dilution. Slides were examined under an epifluorescence microscope, and antibody titers were determined as the reciprocal of the endpoint dilution that showed positive fluorescence.

### 2.9 Transmission-Blocking Assays

The functional cross-reactivity of human anti-*Pvs*48/45-specific IgG against *P. falciparum* parasites was evaluated by SMFA, as described previously [43, 48, 76]. Briefly, 163 µg/mL of the test IgG was mixed with 0.15% - 0.2% stage V gametocytemia of *P. falciparum* NF54 strain and then fed to 3-6 day/old female *Anopheles stephensi* in the presence of human complement. The mosquitoes (N=20 per sample) were maintained for 8 days and dissected to count the number of oocysts in each midgut. A group of mosquitoes (N=40 per sample) were fed with normal human or normal mouse Protein-G purified antibodies and was uses as a negative control.

### 2.10 Statistics

All ELISA data are presented as an average optical density (OD) value from triplicate wells. The Mann-Whitney test was utilized for comparing two groups, and a Kruskal-Wallis test, followed by Dunn’s multiple comparison test, was used for the comparison of more than two groups. A Fischer exact test was used to compare the relative proportion of responding sera between two groups, and a chi-square test was performed for more than two groups. If the chi-square test shows a significant difference among groups, Fischer’s exact test was used to compare two groups at a time, and Bonferroni corrected p-values were calculated. GraphPad Prism software, version 5.0, was used for the analysis. A descriptive statistical analysis of median OD, quartile 1 (Q1), and Q3 was used to measure the variation in OD values. The percent reduction of the mean oocyst intensity (TRA) was calculated using the formula: [(Xc − Xa)/Xc)] × 100, where X is the arithmetic mean oocyst intensity in control (c) and test (a) IgG. The 95% confidence interval and *p*-value for TRA (either from a single assay or two assays) were calculated using a zero-inflated negative binomial model as previously described [76].

## 3. RESULTS

### 3.1 Protein purity and cross-reactivity of r*Pvs*48/45 protein with *P. falciparum* adult immune sera from the four African country donors

After CHO-r*Pvs*48/45 expression, the protein was purified by affinity chromatography, and purity was confirmed by MS/MS [55]. The sequences alignment of full-length *Pvs*48/45 and *Pfs*48/45 proteins confirmed an overall homology of (60.8%) [55], with an even higher homology (> 80%) in the carboxyl region (aa 284-428) (Supl. figure). Sera samples from BF (adults), Mali (adults and children), TZ (adults and children), and from NIG (adults) were analyzed by ELISA for their recognition of the CHO-r*Pvs*48/45. Analysis of adult’s samples indicated a high recognition of CHO-r*Pv*s48/45 proteins in individual sera across the four participant countries, with a proportion of positive responders ranging from 40-94%. Indeed, the responder’s proportion was high in TZ (94%), Mali (90%), and BF (90%), whereas in NIG, the responder’s proportion was lower (40%) (p <0.0001; **Figure 1A**). Furthermore, significant variations were observed in OD values among the different countries (p <0.0001; **Figure 1B**). We observed that OD values were more similar between adult groups from Mali (with a median OD (Q1; Q3) of 0.310 (0.170; 0.450)) and TZ (with a median (Q; Q3) of 0.252 (0.183; 0.342), but these were slightly higher than BF (with a median OD of 0.205 (0.169; 0.302)) which in turn, was significantly higher (p <0.0001) than NIG with a median OD of 0.045 (0.033; 0.086).

**Figure 1:**
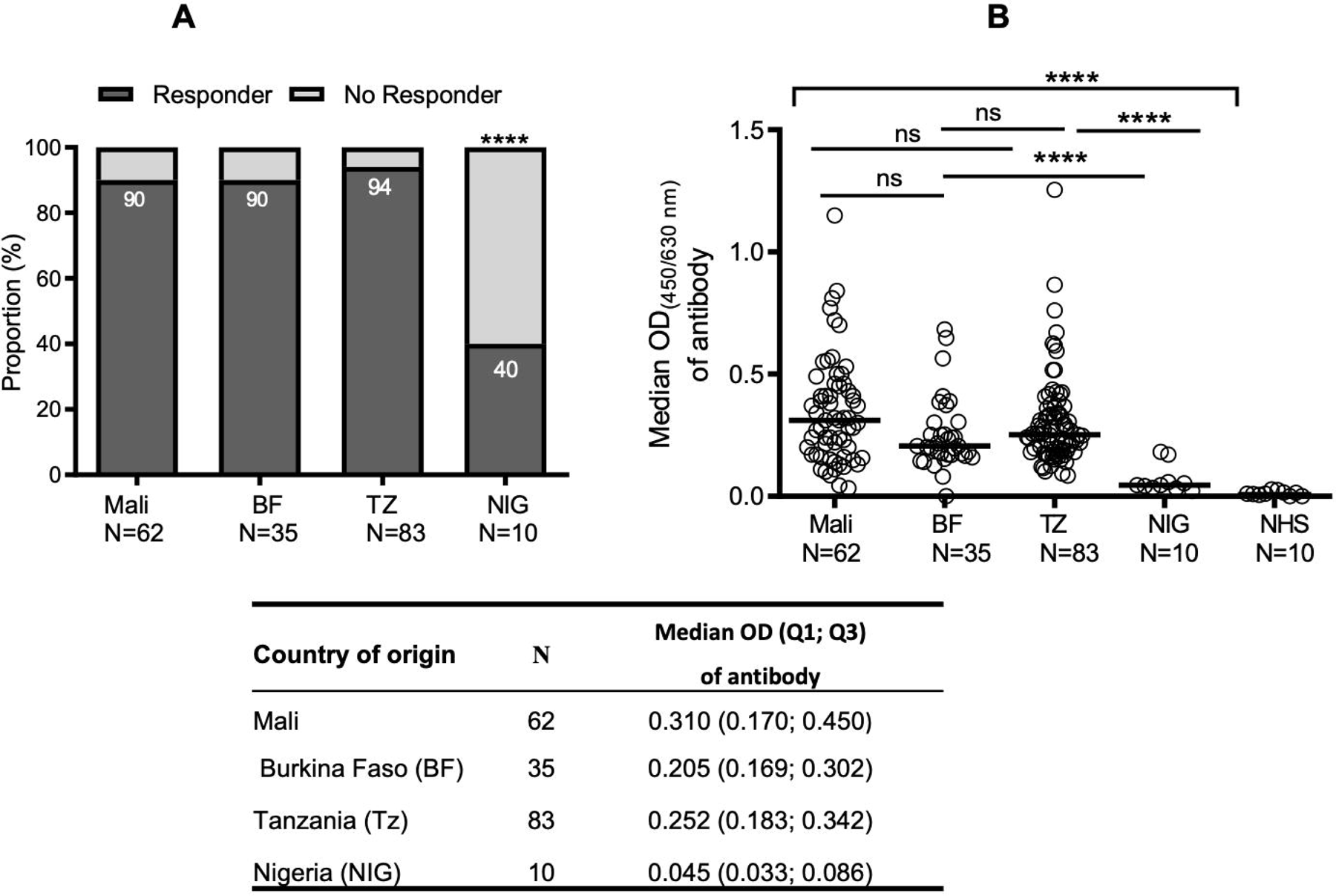
Distribution of cross-reactive antibody responses against r*Pv*s48/45 in P. *falciparum naturally exposed* populations from different African endemic areas. Adult sera from Mali, Burkina Faso (BF), Tanzania (TZ) and Nigeria (NIG) were collected and tested by ELISA against r*Pvs*48/45 at dilution of 1/200. Naïve human sera (NHS) from Swiss donors were used as a negative control. **A**) The proportion of responder samples among Mali, BF and TZ were comparable but significantly higher as compared to NIG. **B)** Global analysis of samples (responder and non-responder) shows that antibody levels (median OD shown as a horizontal black line in the dot plots) for r*Pv*s48/45 were similar among Mali, TZ and BF but significantly higher than NIG. The table shows the median OD, quartile 1 (Q1) and quartile 3 (Q3) of antibody responses. Chi-square test was used to compare responder proportions (A) and Kruskal-Wallis test, followed by Dunn’s multiple comparison test was applied to compare OD values (B). ****p≤ 0.0001; ns; not significant; n, number of sera donors .

### 3.2 Cross-reactivity of r*Pvs*48/45 protein with *P. falciparum* African immune sera with regard to the age and gender

Moreover, we analyzed the cross-reactivity of r*Pvs*48/45 against the individual sera with regards to the age of donors (adult *vs*. children) in Mali and TZ. In Mali, adults demonstrated a significantly higher responder proportion (p <0.0001) to r*Pvs*48/45 and showed a drastically higher median OD of 0.310 (0.170; 0.450) than children (0.151 (0.100; 0.245) (**Figure 2A** and B). For individual sera from TZ, adults and children presented comparable responder rates (p> 0.05; **Figure 2A**) and comparable antibody levels with median OD of 0.252 (0.183; 0.342) and 0.224 (0.176; 0.305) (p <0.05; **Figure 2B**), respectively. However, sera from children in TZ showed a significantly higher responder proportion (p <0.001) and higher OD (p <0.0001) than those from Mali (**Figures 2A** and **B**) while the responder proportion and antibody levels remained comparable in adult donors between the two countries. Of the 97 samples collected from Malian children, gender information was available only for 41 samples, thus we evaluated the gender effect for them. No significant difference was observed in the antibody responder proportion between male (75%; N=16) and female (80%; N=25) young children from Mali (**Figure 3A**), nor in the antibody level between these two groups (**Figure 3B**).

**Figure 2:**
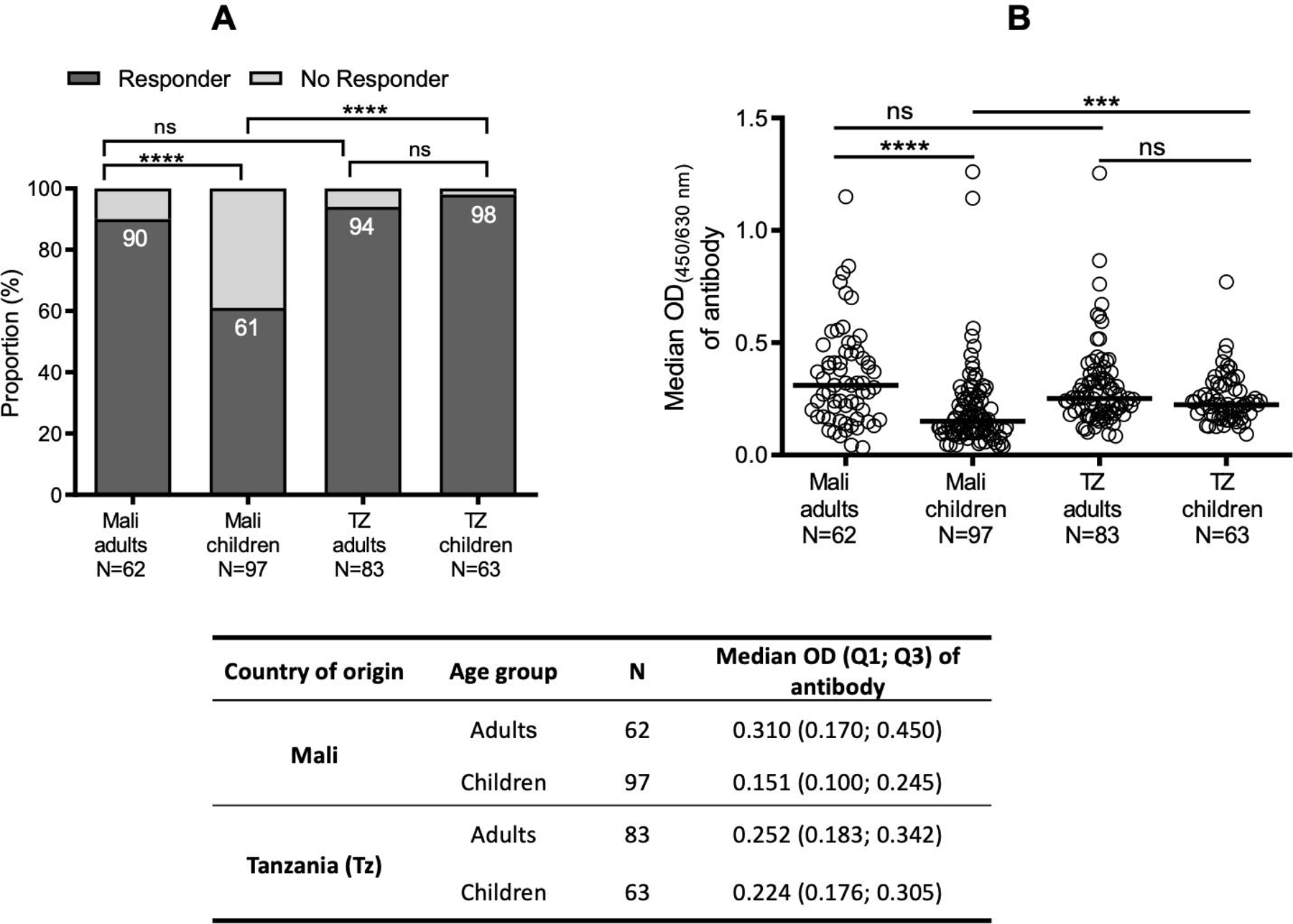
Distribution of cross-reactive antibodies against r*P*vs48/45 protein in adults and children from Mali and Tanzania. In addition to adult samples from Mali and TZ as described in Figure 1, ELISA also tested additional sera from young children (≤ 5 years) for r*Pvs*48/45 protein recognition at the same dilution of 1:200. **A)** The proportion of responders against r*P*vs48/45 was significantly higher for adults than children in Mali, whereas adults and children from TZ showed similar levels. However, children from TZ were more likely to be responders than those from Mali. **B)** Levels of cross-reactive antibodies (median OD shown as a horizontal black line in the dot plots) between adults and children varied significantly in Mali, while antibody levels remained similar between the two age groups in TZ, and between adults from the two countries. The table shows the median, quartile 1 (Q1), and quartile 3 (Q3) OD’s of antibodies against r*Pvs*48/45 protein. Chi-square test was used to compare responder proportions (A) and Kruskal-Wallis test, followed by Dunn’s multiple comparison test was applied to compare OD values (B). ***p≤ 0.001; ****p≤ 0.0001; ns; not significant; N, number of donors.

**Figure 3:**
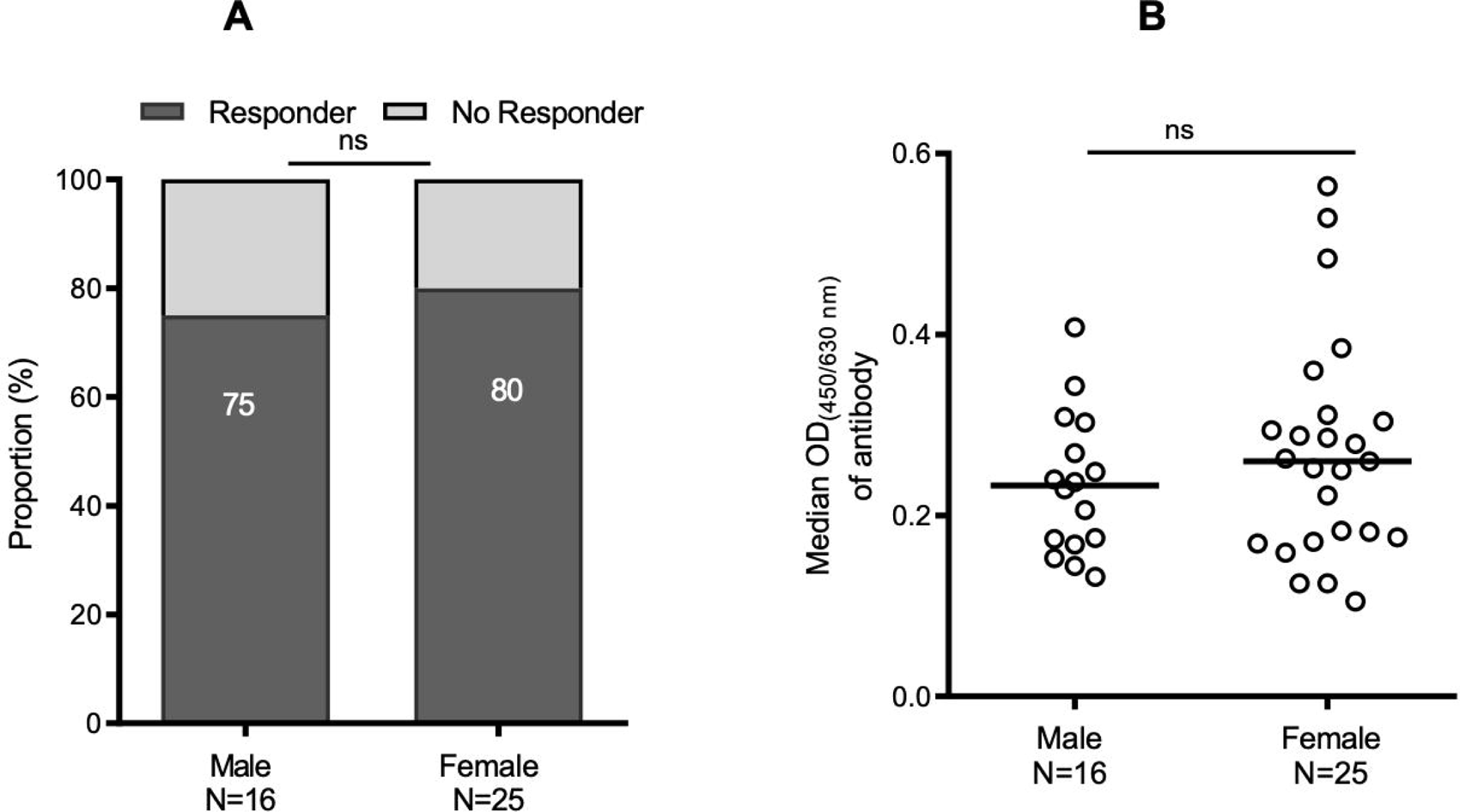
Distribution of cross-reactive antibody responses against r*Pvs*48/45 with regards to the gender of young children in Mali. Cross-reactive antibody responses for r*Pvs*48/45 in young children (≤ 5-year-old) from Mali was further analyzed according to gender (male and female). **A)** Proportion of responder samples for r*Pvs*48/45 remained similar between the male and female. **B)** Cross-reactive antibody levels (median OD shown as a horizontal black line in the dot plots) remained also comparable between male and female children. Fisher’s exact test was performed to compare the proportion of responders between the two gender groups (A) and the Mann-Whitney test was applied to compare the variation of OD’s between male and female individuals (B). N, number of male or female; ns: not significant.

### 3.3 Cross-reactivity of r*Pvs*48/45 protein with African immune sera of *P. falciparum* in competitive ELISA and specific antibody purification

To further characterize the cross-reactive binding of antibodies toward r*Pvs*48/45, the r*Pvs*48/45 protein was inhibited with itself (**Figure 4A**) in competition ELISA in the presence of the two best responder sera from BF adult donors, BF40 and BF70. The competitor r*Pvs*48/45 inhibited binding of anti-r*Pvs*48/45 antibodies to the adsorbed r*Pvs*48/45 protein on the ELISA plate by ∼80% (**Figure 4A**) for both BF40 and BF70 at 300 μg/mL. This suggests that there is no difference or change in conformation of the r*Pvs*48/45 protein between the liquid and the ELISA plate.

**Figure 4:**
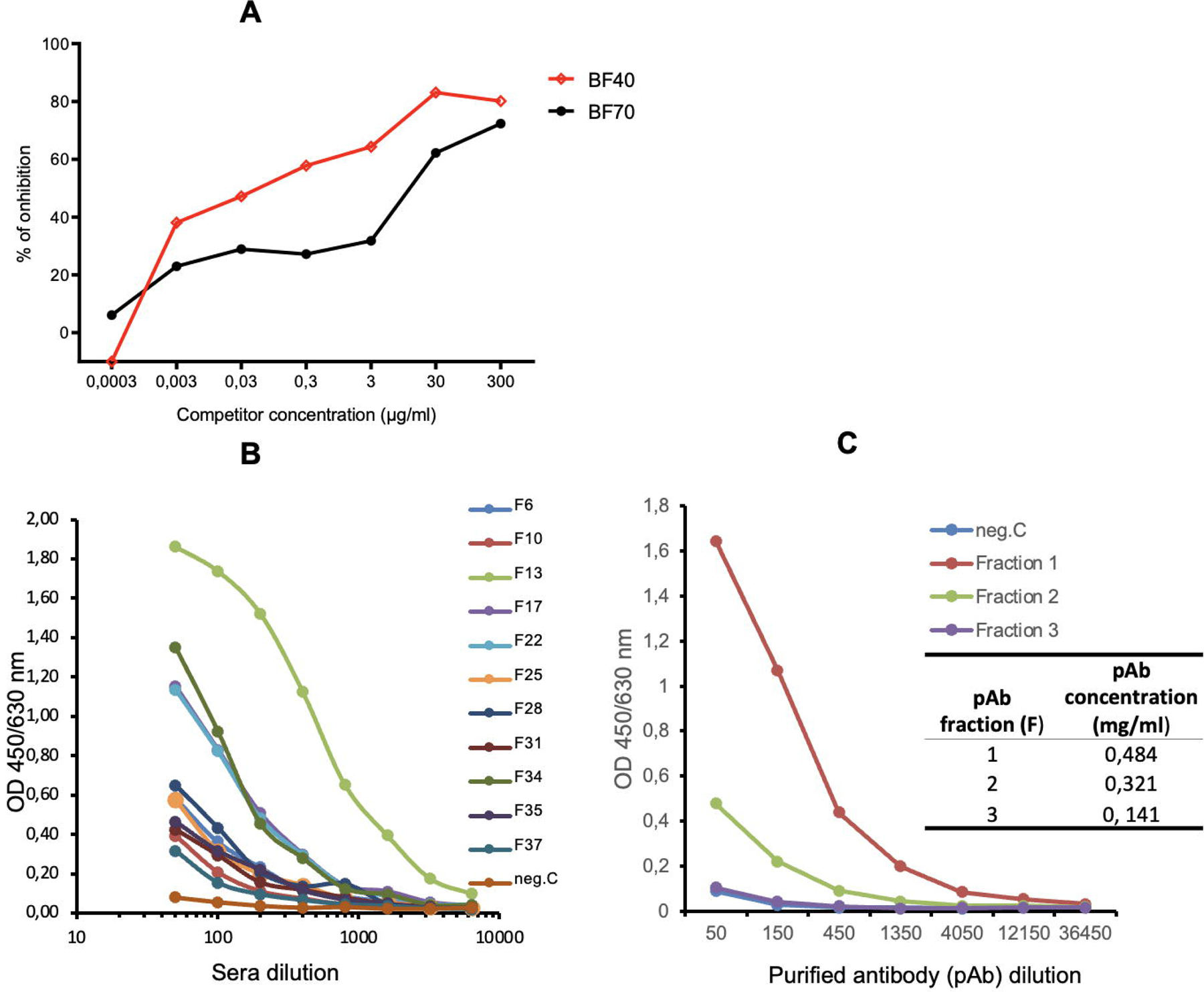
Antibody affinity evaluation and titration curve of cross-reacting sera and purified anti-r*Pvs*48/45 IgG. **A)** Two adult best responder sera from BF (BF40 and BF70) in Figure 1 were used for competition ELISA at dilutions of 1:400 and 1:200, respectively. The 1:400 and 1:200 dilutions gave 50% of the maximum signal in indirect ELISA for each sample. **B)** Adult best responder sera from TZ donors (N=12; F6, F10,…F37) in Figure 1 were tested by ELISA at serial dilutions for their recognition of r*Pvs*48/45. **C)** A pool of the three best responder sera was then used for IgG purification. ELISA (titration curve) studied purified IgG from different eluted fractions (F1, F2 and F3) for r*Pvs*48/45 protein recognition. The inserted table shows the IgG protein concentration in each fraction as measured by nanodrop spectrophotometer. Negative control (neg.C), i.e. naïve human sera, NHS from Swiss naïve donors.

Twelve adult samples from Tanzania (TZ) with the highest response against r*Pvs*48/45 (**Figures** 1 and **2**) were further screened at serial dilutions by ELISA (**Figure 4B**), and three of samples with the strongest responses were then pooled to purify specific IgG against r*Pvs*48/45. Anti-r*Pvs*48/45 responses in each elution fraction from the purification was evaluated by ELISA (**Figure 4C**). As expected, fraction 1 (F1) yielded the highest protein concentration of IgG and the strongest recognition of r*Pvs*48/45 of all the purified fractions and demonstrated a considerably stronger recognition of r*Pvs*48/45 than naive human sera (NHS, as negative control) (**Figure 4C**). Fraction 1 IgG was then later used for the *ex vivo* TB activity assay.

For functional cross-reactivity evaluation, purified r*Pvs*48/45-specific IgG from pooled sera from TZ adults were tested by SMFA against *P. falciparum* parasites. The *ex-vivo* SMFA was carried out using *P. falciparum* gametocytes against fraction F1 of purified IgG (**Table 1**). The purified r*Pvs*48/45-specific IgG showed a 61% inhibition of oocyst intensity (95% CI, 31 to 79 %; p=0.003) at 163 µg/mL (the highest concentration used based on the available IgG).

**Table 1:** Transmission-blocking activity of anti-r*Pvs*48/45-specific IgG from African donors.

| Sample name | 1<sup>st</sup> SMFA |  | 2<sup>nd</sup> SMFA |  | % inhibition | 95% CI<sup>d</sup> | p-value |
| --- | --- | --- | --- | --- | --- | --- | --- |
|  | Mean oocyst | Mosquitoes<sup>c</sup> | Mean oocyst | Mosquitoes<sup>c</sup> |  |  |  |
| *Control IgG<sup>a</sup>* | 9.1 | 36/40 | 15.3 | 31/40 |  |  |  |
| *Anti-rPvs48/45 IgG<sup>b</sup>* | 4.3 | 13/20 | 4.9 | 11/20 | 61.2 | 31.2 to 78.6 | 0.003 |

<sup>a</sup> In the first assay, normal human IgG at 3750 µg/mL was used as a control and normal mouse IgG at 750 µg/mL was used in the second assay
<sup>b</sup> The anti-rPvs48/45-specific IgG from Tanzania adults was tested at 163 µg/mL in both assays
<sup>c</sup> Number of infected mosquitoes (mosqs) / Number of dissected mosquitoes
<sup>d</sup> 95% confidence interval

### 3.4 *Mice immunization with P. falciparum* gametocytes boost anti-*Pvs*48/45 antibody responses which recognize *P. falciparum* gametocytes in IFAT

In ELISA, a vigorous anti-CHO-r*Pvs*484/45 antibody response was observed in BALB/c mice that were immunized with CHO-r*Pvs*484/45 protein (round symbols) on days 0, 20 and 40 (thin arrows) and not seroconversion were observed in the control group (square symbols). The antibody responses of the experimental group declined subsequently, and reached to baseline levels on day 260. When the mice were inoculated with a single dose of *P. falciparum* gametocytes (5 x10^5^), they produced a significant boost of the anti r*Pvs*48/45 ELISA antibody titers (**Figure 5A**). Additionally, IFAT carried out using the pooled sera collected on day 320 (60 after *P. falciparum* boost) showed strong reactivity with enriched *P. falciparum* gametocyte preparation (**Figure 5B**, bottom panel). The pooled sera collected on day 120 (before *P. falciparum* boost) reacted strongly with *P. vivax* homologous antigens (upper panel).

**Figure 5:** Mice immunized with *P. falciparum* gametocytes boost the anti-Pvs48/45 antibody responses which recognized *P. falciparum* gametocytes in IFAT. **A**) Groups of experimental and control mice were immunized on days 0, 20, and 40 (thin arrows) with r*Pvs*48/45 protein (round symbol) or placebo (square symbol), respectively, then boosted with 5 x 10^5^ *P. falciparum* NF-54 gametocytes emulsified in Montanide ISA-51 on day 260 (bold arrow). ELISA determined the antibody titers in individual mice at different time points. Mean and standard deviation in log-transformed antibody titers are shown. **B)** A pooled mouse sera collected on day 120 (before *P. falciparum* boost) were tested by IFAT with *P. vivax* (upper panel), and another pooled sera collected on day 320 (60 after *P. falciparum* boost) were tested with *P. falciparum g*ametocytes (bottom panel). Parasites were seen under light (left) or epifluorescence (right) microscopy with a 100X objective lens are shown. Picture scale 238 μm.

## 4. DISCUSSION

*Pvs*48/45 is a protein expressed on the surface of *P. vivax* gametocytes, known to be involved in parasite fertilization [47, 48, 58]. Both *P. falciparum* and *P. vivax* 48/45 proteins are well-established as targets of natural antibody responses to parasitic sexual stages, which have shown important TB activity in *ex vivo* assays. Consequently, they are currently being pursued as TB vaccines candidates [45, 46, 77–79].

This study indicates that sera from a significant proportion of donors (40-94%) living in *P. falciparum*-endemic areas of Africa cross-recognize the r*Pvs*48/45 protein. The high and consistent recognition of r*Pvs*48/45 by sera from different endemic regions of Africa, with no *P. vivax* transmission at the time of sera collection, is remarkable. Unlike previous studies in endemic areas of Latin America, where both parasites coexist and tend to induce cross-boosting of the antibody responses in natural conditions and in mice immunization [63, 80, 81], the present study clearly demonstrates cross-reactivity of *Pvs*48/45 against samples from *P. falciparum-*endemic areas. This cross-reactivity is most likely explained by the significant amino-acid sequence homology (∼ 60,8%) between *P. vivax* and *P. falciparum Ps*48/45 [57, 59]. This feature makes this protein a promising target for the development of an effective cross-species vaccine.

The communities analyzed in this study have been historically exposed to variable *P. falciparum* transmission intensities, according to the 2018 WHO report [82] . The r*Pvs*48/45 recognition intensity, as determined by the level of specific anti-r*Pvs*48/45 and the percentage of positive responders in each of the four endemic countries, may be correlated with the relative transmission intensity and episodes history of malaria in these countries. Further research is now necessary to test these hypotheses. Furthermore, despite a high proportion of positive serological responses, our research highlights significant differences in antibody levels (OD value) based on demographics. While cross-reactive antibody levels were more comparable and higher for Mali and TZ where samples were collected from rural sites, than those from BF or NIG, collected from urban sites. These findings support the argument that populations living in rural communities and small villages are more likely to be exposed to malaria vectors than those living in metropolitan areas [83].

In addition to location, age played an important role in antibody response. Adults made up a greater proportion of responders and demonstrated higher antibody levels than children, mostly in Mali. This suggests that cross-reactive immune responses to the r*Pvs*48/45 protein may increase with age, which is argued to be a result of adults being exposed for longer periods of time, as a function of their age. Hence, these cross-reactive anti-r*Pvs*48/45 antibodies are likely acquired early in childhood and are then boosted throughout the donors’ life, most likely in response to subsequent *P. falciparum* infections. This age-related increasing trend of the specific immune response in the naturally exposed population has also been demonstrated with other malaria antigens, although such immunity was due to antigens specific to *Plasmodium* species [53, 77, 84–86]. Moreover, recent sero-epidemiological studies, performed with sera from malaria-endemic regions where both *P. vivax* and *P. falciparum* are co-transmitted, suggest that frequent exposure to *P. falciparum* infections results in the maintenance of anti-*P. vivax* antibodies [45, 55, 56, 63, 79]. However, this current study did not determine which Ps48/45 segments were cross-reactive, or demonstrate which conserved protein domains are prone to be restricted despite the recognition of heterologous parasites that was observed [62, 87, 88]. One of the limitations of this study is also that these sera were not tested with the recombinant *Pf*s48/45 protein. Although, in Africa, where *Pf* is present, it has been shown that a naturally acquired antibody response to the recombinant *Pf*s 48/45 protein is clearly detected [89–91]. Another limitation of the study is that the gender effect was assessed only in a part of Malian children samples, where genter information was available. Further study is required whether there is no gender difference in other populations.

Altogether, the consistent reactivity of antibodies against r*Pvs*48/45 protein in both *P. falciparum* and *P. vivax* parasites under natural conditions in distant continents with significant epidemiological differences, as well as in animal models, correlates with the protein sequence conservation. The results of populations naturally exposed to *P. falciparum* are also in agreement with the cross-reactivity and cross-boosting effects observed in mice experimentally immunized with r*Pfs*48/45 and r*Pvs*48/45 *E. coli* recombinant products [55, 63]. Moreover, the anti-r*Pvs*48/45 response appears to be consistent with the high proportion of antibodies against *Pvs*48/45 reported in adults from malaria-endemic areas of Latin America, where malaria transmission is significantly lower [62, 92].

More importantly, the significant *ex-vivo* reduction by 61.2% of *P. falciparum* oocyst development in *An. stephensi* fed with *P. falciparum* gametocytes by affinity-purified anti-r*Pvs*48/45 IgG is of interest and encourages investing further efforts into characterizing the functional domains to model multispecies TB vaccine development. Although the cross-species *ex vivo* TB activity is suboptimal, the likelihood of inducing robust TB in both species through vaccination is highly likely.

The recognition of the native proteins of the two species in IFAT assays, and the *P. falciparum ex-vivo* TB activity and the *P. falciparum* gametocyte boosting of anti-CHO r*Pvs*48/45 antibodies in mice, represent solid foundations that support r*Pvs*48/45 as a target for a TB vaccine. Moreover, the current epidemiological data, together with the ELISA cross-species reactivity and the TB capacity of r*Pvs*48/45-specific antibodies purified from *P. falciparum* semi-immune individuals against, support the further development of a cross-species TB vaccine.

## Supporting information

Supplemental figure 2: Recombinant CHO-rPvs48/45 protein analysis in western Blott

Supplemental figure 1. Sequence homology between the Pvs48/45 and Pfs48/45 proteins

## 6. Author Contributions

SB, GP, SH and MAH designed the experiment. SB, KM, IA, DK, NCI, VA, SMH and CL performed most experiments, tests, and analyses. SB, KM, GP, SMH, SH and MAH wrote the manuscript. MAG, SAD, SO, IN, AVK and MD contributed to antigen and sample processing, and manuscript revisions. All authors read and approved the submitted version.

## Acknowledgments

We are grateful for the participation of the community from malaria-endemic countries of Mali, Tanzania, Burkina Faso and Nigeria, as well as Swiss volunteers. We would like to thank Drs Marcel Tanner and Ingrid Felger for sharing sera from malaria endemic areas and acknowledge the intramural program of National Institute of Allergy and Infectious Disease for the valuable contribution with *P. falciparum* functional assays.

## 8. Data Availability Statement

The raw data supporting the conclusions of this article will be made available by the authors, without undue reservation.

