## Supplementary figures and images for "Cross-reactivity of r*Pvs*48/45, a recombinant *Plasmodium vivax* protein, with sera from *Plasmodium falciparum* endemic areas of Africa"

### Supplemental figure 2: Recombinant CHO-rPvs48/45 protein analysis in western Blott

#### Slide 1
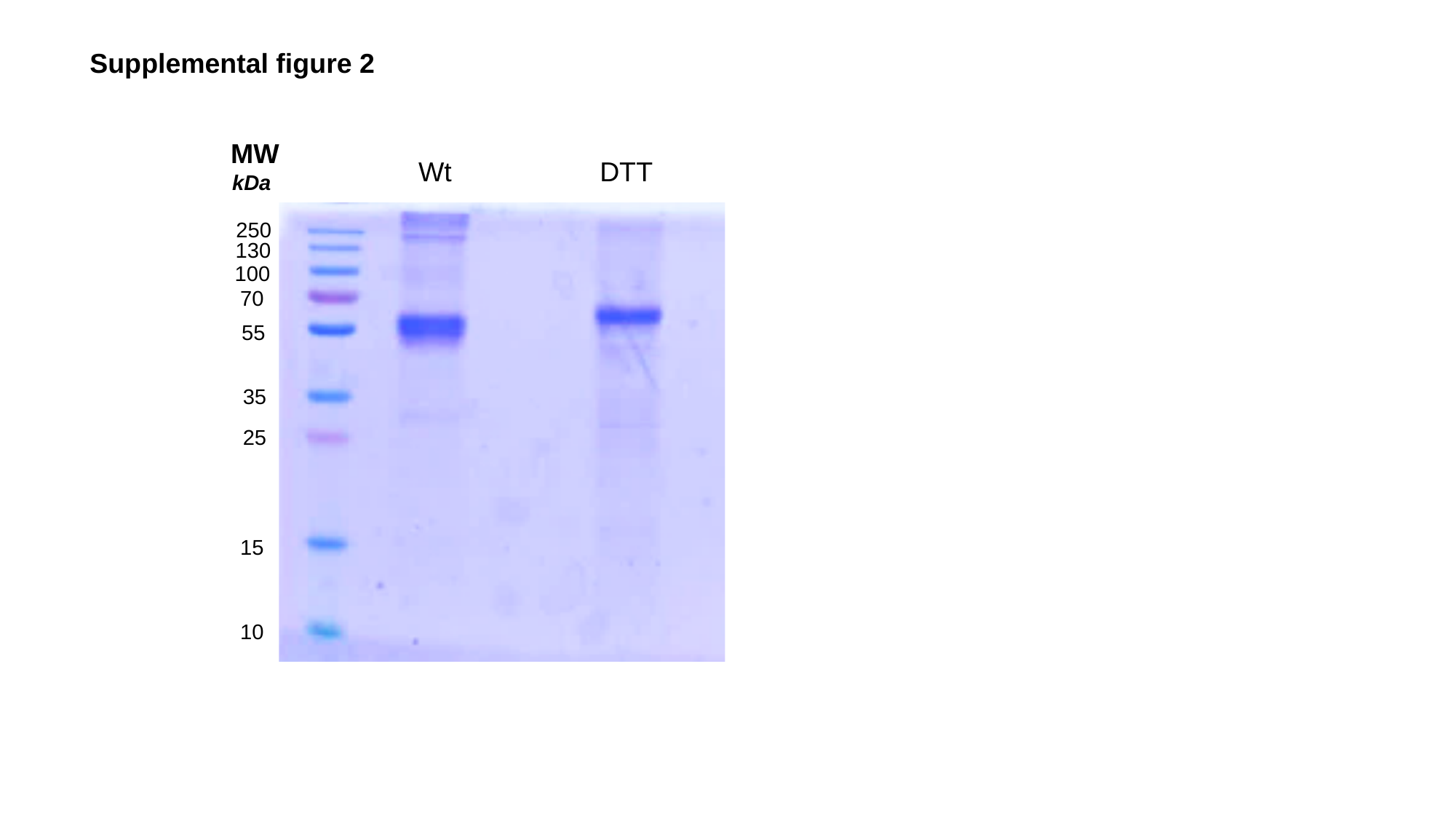

Supplemental figure 2
MW
kDa
250
130
100
70
55
35
25
15
10
Wt
DTT
