## Supplemental figure 1. Sequence homology between the Pvs48/45 and Pfs48/45 proteins for "Cross-reactivity of r*Pvs*48/45, a recombinant *Plasmodium vivax* protein, with sera from *Plasmodium falciparum* endemic areas of Africa"

### Slide 1
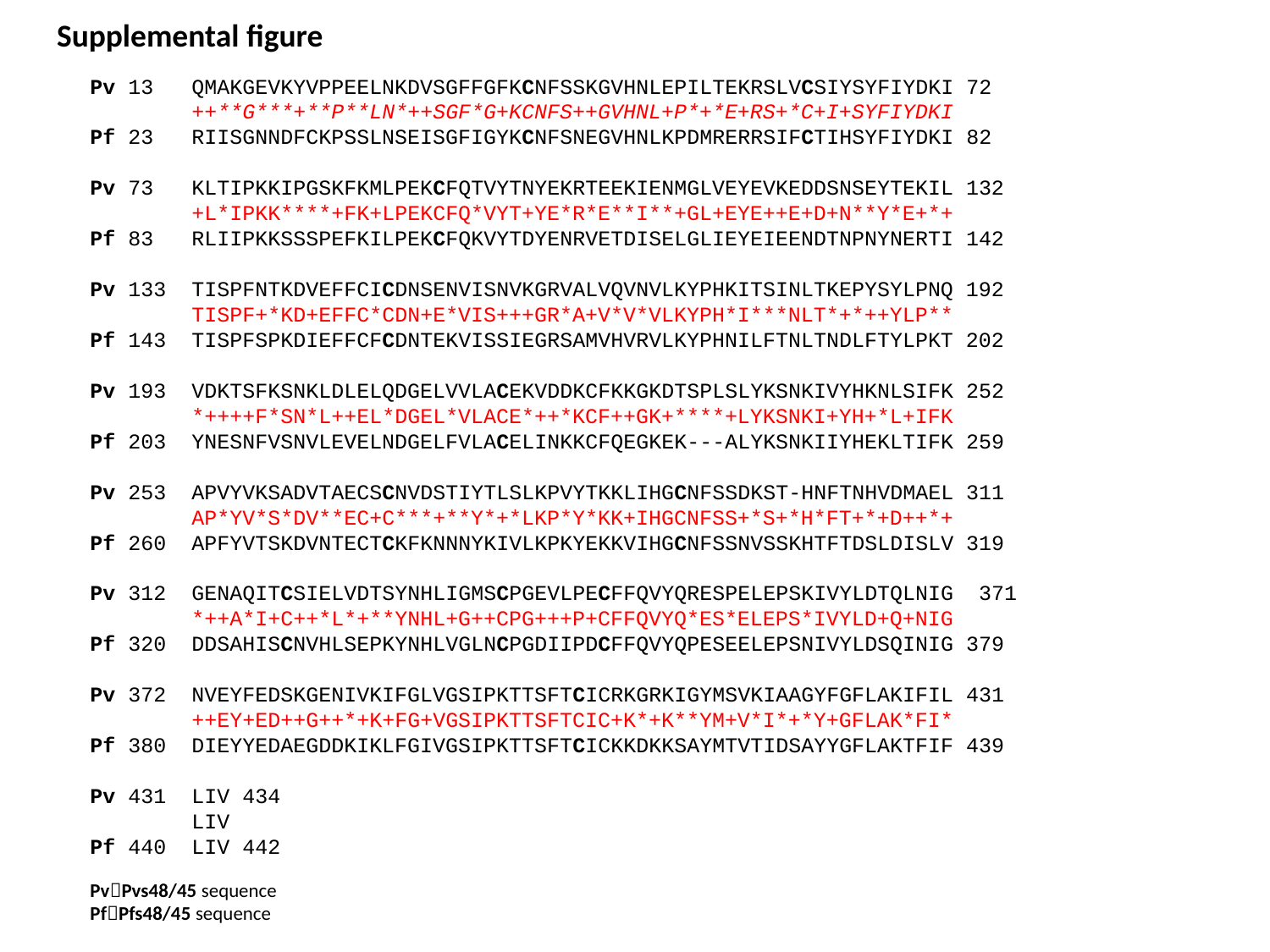

Supplemental figure
Pv 13 QMAKGEVKYVPPEELNKDVSGFFGFKCNFSSKGVHNLEPILTEKRSLVCSIYSYFIYDKI 72
 ++**G***+**P**LN*++SGF*G+KCNFS++GVHNL+P*+*E+RS+*C+I+SYFIYDKI
Pf 23 RIISGNNDFCKPSSLNSEISGFIGYKCNFSNEGVHNLKPDMRERRSIFCTIHSYFIYDKI 82
Pv 73 KLTIPKKIPGSKFKMLPEKCFQTVYTNYEKRTEEKIENMGLVEYEVKEDDSNSEYTEKIL 132
 +L*IPKK****+FK+LPEKCFQ*VYT+YE*R*E**I**+GL+EYE++E+D+N**Y*E+*+
Pf 83 RLIIPKKSSSPEFKILPEKCFQKVYTDYENRVETDISELGLIEYEIEENDTNPNYNERTI 142
Pv 133 TISPFNTKDVEFFCICDNSENVISNVKGRVALVQVNVLKYPHKITSINLTKEPYSYLPNQ 192
 TISPF+*KD+EFFC*CDN+E*VIS+++GR*A+V*V*VLKYPH*I***NLT*+*++YLP**
Pf 143 TISPFSPKDIEFFCFCDNTEKVISSIEGRSAMVHVRVLKYPHNILFTNLTNDLFTYLPKT 202
Pv 193 VDKTSFKSNKLDLELQDGELVVLACEKVDDKCFKKGKDTSPLSLYKSNKIVYHKNLSIFK 252
 *++++F*SN*L++EL*DGEL*VLACE*++*KCF++GK+****+LYKSNKI+YH+*L+IFK
Pf 203 YNESNFVSNVLEVELNDGELFVLACELINKKCFQEGKEK---ALYKSNKIIYHEKLTIFK 259
Pv 253 APVYVKSADVTAECSCNVDSTIYTLSLKPVYTKKLIHGCNFSSDKST-HNFTNHVDMAEL 311
 AP*YV*S*DV**EC+C***+**Y*+*LKP*Y*KK+IHGCNFSS+*S+*H*FT+*+D++*+
Pf 260 APFYVTSKDVNTECTCKFKNNNYKIVLKPKYEKKVIHGCNFSSNVSSKHTFTDSLDISLV 319
Pv 312 GENAQITCSIELVDTSYNHLIGMSCPGEVLPECFFQVYQRESPELEPSKIVYLDTQLNIG 371
 *++A*I+C++*L*+**YNHL+G++CPG+++P+CFFQVYQ*ES*ELEPS*IVYLD+Q+NIG
Pf 320 DDSAHISCNVHLSEPKYNHLVGLNCPGDIIPDCFFQVYQPESEELEPSNIVYLDSQINIG 379
Pv 372 NVEYFEDSKGENIVKIFGLVGSIPKTTSFTCICRKGRKIGYMSVKIAAGYFGFLAKIFIL 431
 ++EY+ED++G++*+K+FG+VGSIPKTTSFTCIC+K*+K**YM+V*I*+*Y+GFLAK*FI*
Pf 380 DIEYYEDAEGDDKIKLFGIVGSIPKTTSFTCICKKDKKSAYMTVTIDSAYYGFLAKTFIF 439
Pv 431 LIV 434
 LIV
Pf 440 LIV 442
Pv🡪Pvs48/45 sequence
Pf🡪Pfs48/45 sequence
